# The paternally derived genome opposes seed dormancy induction by the mother plant in Arabidopsis

**DOI:** 10.1101/2024.11.07.622415

**Authors:** Thiago Barros-Galvão, Xiaochao Chen, Steven Penfield

## Abstract

Seed dormancy in Arabidopsis is known to be mediated by the interaction of maternal and zygotic genomes during seed maturation. While studies have revealed the extensive influence of maternal processes on dormancy and germination, less is known about the influence of the father. Here we exploit differences in ploidy to explore the role of the paternal genome on progeny seed dormancy. We show that paternal genome acts to reduce seed dormancy regardless of maternal genome dose, resulting in lower dormancy in tetraploid Arabidopsis versus genetically identical diploids. We show that this paternal effect requires synthesis of RNA Polymerase IV-dependent RNAs in the male gametophyte which oppose the dormancy-inducing effects of maternal siRNAs on seed coat and endosperm development. We conclude that the paternal genome has evolved to subvert the dormancy-inducing role of the mother plant in progeny seeds.

## Introduction

Seed dormancy is under substantial control of the mother plant (Penfield, 2017). In Arabidopsis this includes the maternal seed coat because the endosperm cuticle is necessary for dormancy and is synthesised by the inner integument (Loubéry et al., 2018). Seed dormancy is highly dependent on the seed production environment and environmental effects on seed dormancy are mediated by ABA transport from the mother plant and frequently accompanied by morphological and biochemical changes to maternal fruit and seed coat tissues in addition to the physiology of the zygote (Chen et al., 2024; Chandler et al., 2024). Response to environmental signals by developing zygotes is also influenced by the epigenetic state of maternally-derived genome in the endosperm (Iwasaki et al., 2019; Sato et al., 2021; Chen et al., 2023), underlining the multi-faceted nature of the maternal influence on progeny seed behaviour.

The maternal regulation of seed dormancy is integral to bet-hedging germination strategies in which the mother plant creates cohorts of seeds with varying dispersal strategies, exposing individual progeny seeds to varying risk profiles designed to benefit the fitness of the mother plant alone in an example of parent-offspring conflict (Penfield, 2017). This explains the phenomenon of seed dimorphism whereby mothers produce a fixed or varying ratios of seeds of different sizes and dormancy to fulfil different dispersal niches: a good example in the Brassicaceae is the genus *Cakile* (Maun and Payne, 1989). Because fathers may only parent of a fraction of a seed lot, parental conflict theory predicts benefits to fathers that can ensure increased resourcing of individual progeny or confer other competitive advantages such as fast germination and strong establishment. In interploidy crosses of Arabidopsis it has been hypothesised that this conflict manifests as variation in seed size, driven by parentally derived genome dose in the endosperm and dependent on DNA methylation (Scott et al., 1998; Xiao et al., 2006).

Polyploidy is common in plant evolution and may confer advantages over diploids relating to seedling size driven by the increase in genome content. In the angiosperm endosperm the mother has a genome dosage advantage and mother plants can detect and initiate abortion of seeds pollinated by fathers with excess ploidy through the process known as ‘triploid block’. Triploid block is dependent on the action of RNA Polymerase IV (Pol IV) -mediated siRNA production in the parent plants (Erdmann et al., 2017; Martinez et al., 2018; Grover et al., 2018). Polyploidy can also influence fitness via seed dormancy and germination traits such as germination speed or establishment vigour (Eliasova and Munzbergova, 2014). However, making general empirical determinations of the effect of polyploidisation on seed dormancy using wild species is problematic because of unknown genetic differences between field-collected diploids and polyploids and rapid selection for dormancy-associated traits (Huang et al., 2010).

Here we show that paternal genome dose has strong dormancy-reducing effects that are only partially offset by maternal genome dose, leading to strong dormancy reductions in polyploid Arabidopsis compared to genetically similar diploid progenitors. We show that just as in the case of triploid block this process is dependent on parental Pol IV-derived siRNAs and associated with changes to seed coat tissues in addition to the endosperm.

## Results

Loss of *vel3* in in the central cell leads to low seed dormancy and hyperactivation of senescence-promoting pathways in the endosperm during seed maturation, as well as activation of seed abortion via the so-called ‘triploid block’ (Chen et al., 2023). In Arabidopsis the Landsberg erecta (Ler) ecotype facilities bypass of the triploid block due to loss of MIR845 and paternal siRNA production by RNA polymerase IV (Erdmann et al., 2017). To test whether activation of the triploid block is essential for the dormancy effect we backcrossed *vel3-1* into Ler and found that in the Ler background *vel3-1* mutants have normal seed dormancy when set at 15°C (Figure 1A). Furthermore, crossing *vel3-1* (Ler) mothers to *vel3-1* (Col) fathers resulted in seeds with a wide range of dormancy states but frequently showed low dormancy, suggesting that the paternal Ler genotype plays a role in the rescue of the dormancy phenotype. This was accompanied by a small increase in the frequency of aborted seeds (Figure 1B).

**Figure 1.**
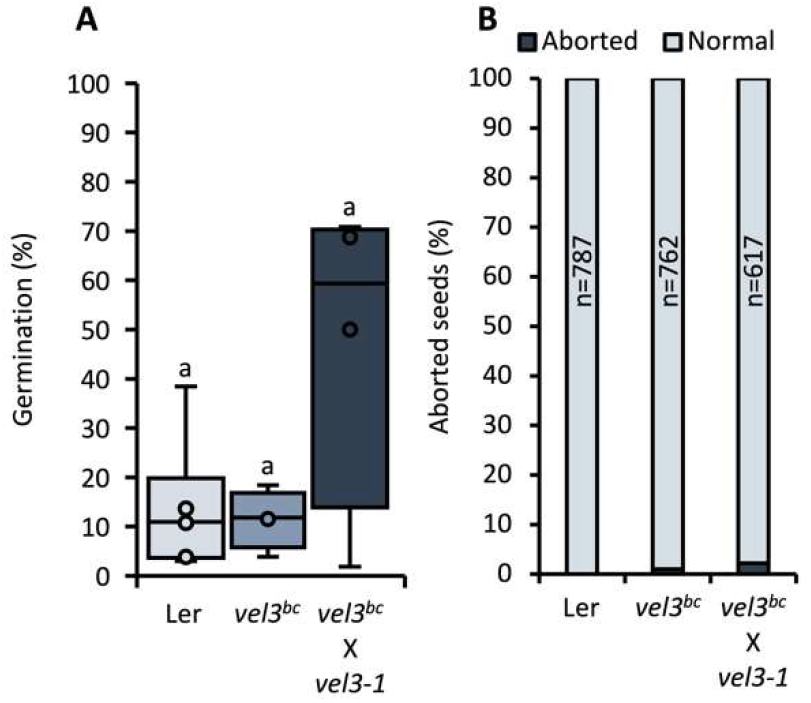
VEL3 does not affect seed dormancy and abortion in Landsberg *erecta* background. **A**. Germination of Ler, *vel3-1* introgressed in Ler (*vel3*^bc^), and maternal *vel3*^bc^ crossed with paternal *vel3-1* in Col seeds set at 15 °C. Significant differences are shown using ANOVA with Tukey post hoc test (P > 0.05 n = 4-6). **B**. Seed abortion frequency (%) in Ler, *vel3-1*^bc^, and *vel3-1*^bc^ X *vel3-1*. Total number of seeds analyzed for each genotype are shown.

The previous result suggested that dormancy-regulation is intrinsically linked to activation of triploid block and observable in seeds which do not fully complete the abortion process. This lead us to hypothesise that changes in parental genome dose should affect seed dormancy in backgrounds that generate sufficient viable seeds. To test this we exploited the ability of the Ler accession to evade triploid block generating seeds from interploidy crosses and measuring dormancy by setting seeds at low temperatures (Figure 2A). This showed that seeds with paternal excess exhibited a strong low dormancy phenotype, whereas seeds with maternal excess were marginally more dormant than WT. However, more surprisingly, the dormancy phenotype of paternal genome excess was shared with tetraploid control seeds, suggesting that the effect of paternal genome dose is to some degree independent of the activity of the maternal genotype. To confirm this finding we conducted two further experiments: firstly, we set seed from Col-0 diploid and tetraploid plants and found again that tetraploid seed has lower dormancy (Figure 2B). Finally we exploited the ability of the *Jason-1* (*jas-1)* mutant to produce unreduced gametes at high frequency comparing the dormancy of seeds set from 1n and 2n pollen on the same mother plant (Figure 2C). Again we found that pollination of diploid mothers with 2n pollen resulted in high germination rates, but also that pollination of tetraploid mothers with 2n pollen led to a similar outcome. However some evidence of a balancing role for the maternal genome was found because tetraploid mothers pollinated with 1n pollen produced seeds with more dormancy than wild type. However, taken together our data show that increasing paternal genome dose results in lower dormancy and while this effect may be moderated by maternal genome dose, increasing maternal genome dose is insufficient to restore a WT phenotype.

**Figure 2.**
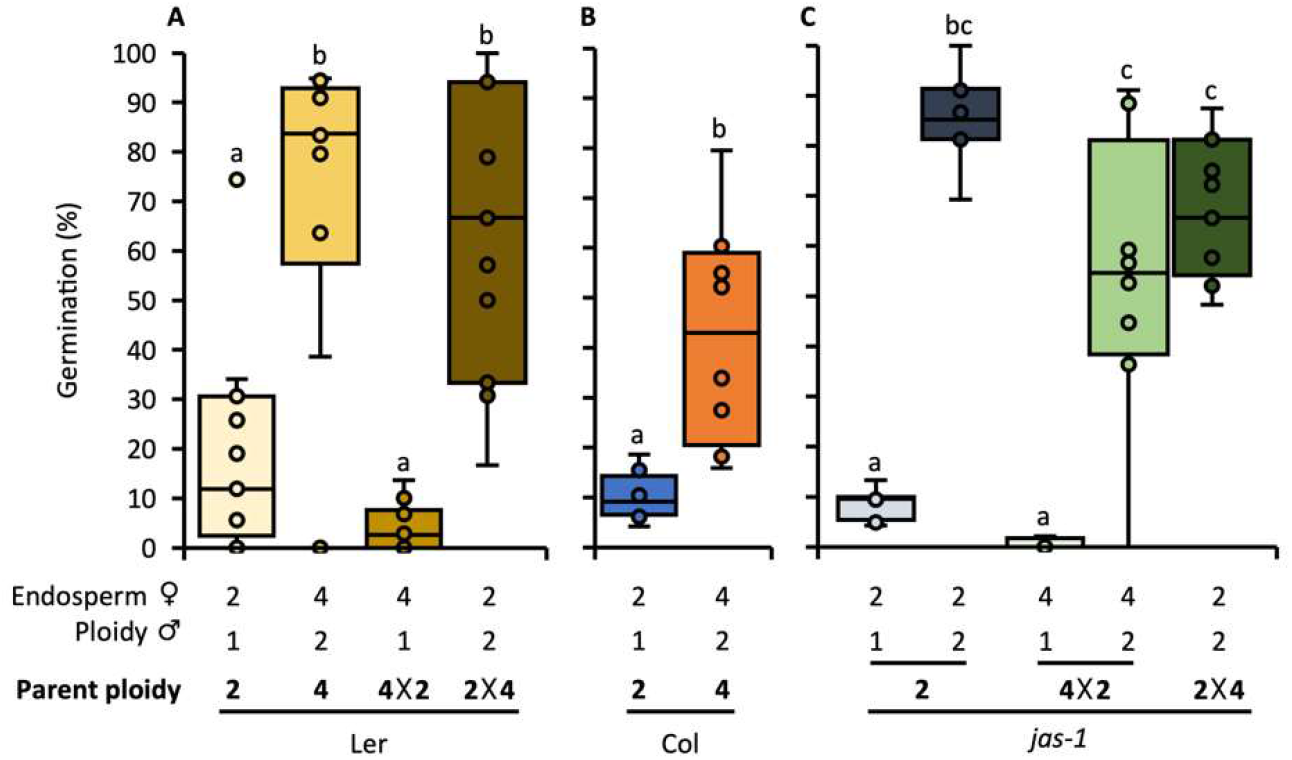
Seed dormancy is reduced by the increases in paternal genome dose. **A**. Germination of freshly harvested L*er* diploid (2n), L*er* tetraploid (4n), maternal Ler tetraploid crossed with paternal Ler diploid (4×2), and reciprocal crosses set at 15 °C. Significant differences are shown using ANOVA with Tukey post hoc test (P < 0.05 n = 8). **B**. Germination of freshly harvested Col-0 2n and 4n seeds set at 15 °C. Significant differences are shown using ANOVA with Tukey post hoc test (P < 0.05 n= 8). **C**. Germination of freshly-harvested *jas-1* diploid seeds (from diploid *jas-1* plants), *jas-1* triploid seeds (from diploid *jas-1* plants), *jas-1* triploid (from maternal tetraploid *jas-1* crossed with paternal diploid *jas-1*), and seeds from reciprocal crosses between *jas-1* 2n and 4n plants set at 15 °C. Significant differences are shown using ANOVA with Tukey post hoc test (P < 0.01 n = 8-11).

Seed dormancy in Arabidopsis depends primarily on the dormancy-promoting activity of the phytohormone abscisic acid (ABA) which accumulates in seeds during seed development and maturation (Karssen et al., 1983; Koornneef et al., 1989). Because the *jas-1* mutant enables us to compare seeds with 3n and 4n endosperms of the same mother plants we measured ABA in seeds with 3n and 4n endosperms (Figure 3A). ABA levels were identical on a per gram basis, strongly indicating that the concentration of ABA is not afected by ploidy. We also tested ABA sensitivity of germination of *jas-1* seeds and diploid and tetraploid seeds. Diploid and tetraploid seeds displayed identical ABA sensitivity, whereas triploid progeny of *jas-1* mutants had marginally higher germination (Figure 3B). Taken together, changes to ABA sensitivity do not account for the dormancy phenotypes observed.

**Figure 3.**
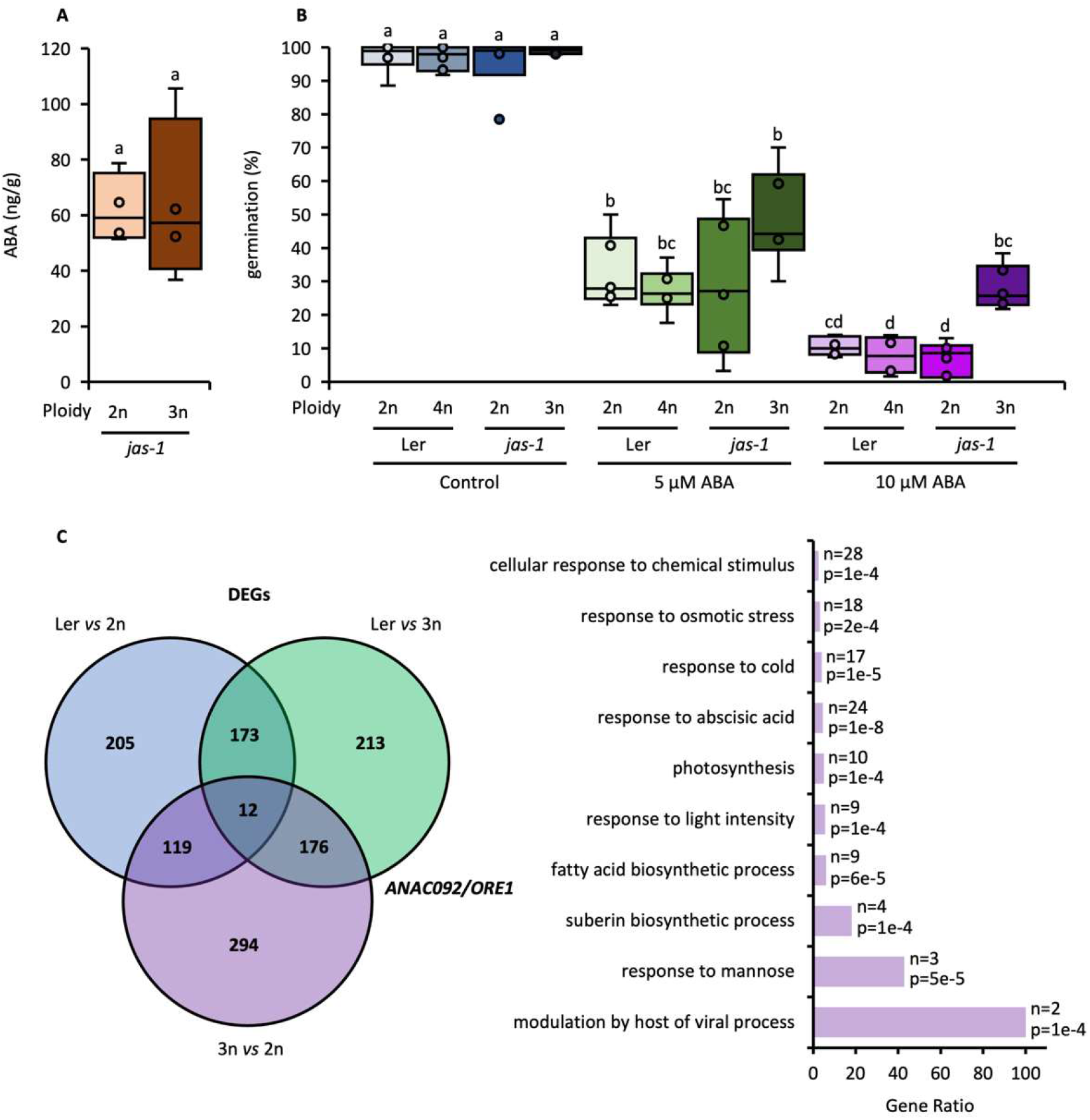
ABA levels and sensitivity play no role in the dormancy phenotypes observed in polyploids. **A**. ABA levels measured by LC-MS in dry seeds of *jas-1* diploid (2n) and *jas-1* triploid (3n), from diploid *jas-1* plants, set at 15 °C. Significant differences are shown using ANOVA with Tukey post hoc test (P > 0.05 n = 4). **B**. Germination of Col-0, Ler and *jas-1* seeds with the indicated embryo ploidy levels set at 22 °C in the presence or absence of exogenous ABA. Significant differences are shown using ANOVA with Tukey post hoc test (P < 0.05 n = 6). **C**. Comparison of mis-regulated genes in Ler, diploid (2n) *jas-1*, and triploid (3n) *jas-1* in the mature endosperm, gene in bold (*ANAC092*/*ORESARA1*; *ORE1*) plays a key role in the dormancy phenotype of *vel3-1* seeds (Chen et al., 2023). GO-Term analysis of up-regulated genes in endosperm of 3n vs 2n mature seeds set at 15 °C, showing gene function category, p-values and number of genes in each category.

Next we compared mature endosperm transcriptomes of *jas-1* diploid seeds to those of *jas-1* triploid seeds and WT Ler. We found around 188 genes mis-regulated in mature endosperms common to *jas-1* 3n seeds vs Ler and jas-1 3n seeds versus *jas-1* 2n seeds (Figure 3C). Genes up-regulated in *jas-1* 3n seeds were counter-intuitively enriched in genes included in GO Term response to ABA: these included the regulator of ABA-induced senescence *ANAC092*/*ORE1*, the up-regulation of which is also important for the low dormancy of *vel3-1* seeds (Chen et al., 2023).

This shows that the likely mechanism of dormancy reduction in seeds with increased paternal genome dose is likely identical to that previously identified in *vel3-1*.

Paternal siRNAs are required for manifestation of the triploid block and are generated in the vegetative cell of the male gametophyte by the action of RNA polymerase IV and act by opposing the effects of maternal siRNA in the endosperm (Borges et al., 2018; Martinez et al., 2018). Furthermore, loss of RNA PolIV-mediated siRNA production results in low dormancy when seeds are set at low temperatures (Iwasaki et al., 2019), but it remains unclear whether this effect is uniparental. To test whether parental RNA PolIV is required for dormancy we compared dormancy in *nrpd1* mutants, WT and reciprocal crosses (Figure 4A). We confirmed that homozygous *nrpd1* mutants have a low dormancy phenotype and further could show that this is conferred primarily by loss of maternal siRNAs. Uniparental loss of *nrpd1* also resulted in seed size increases, while maternal loss alone also gave rise to seeds with a paler seed coat (Figure 4B). Therefore maternal siRNAs are important for dormancy imposition. To understand the interaction of maternal and paternal siRNAs we analysed dormancy in *jas-1 nrpd1* double mutants (Figure 4B), comparing the dormancy of double mutant seeds with 3n endosperms and 4n paternal excess endosperms (Figure 4C). This is general showed that maternal *nrpd1* and paternal *nrpd1* have antagonistic effects on seed dormancy. Loss of maternal nrpd1 caused low dormancy regardless of paternal genome dose whereas paternal loss also abolished the reduced dormancy of *jas-1* triploid seeds and increased dormancy compared by bi-parental loss of *jas-1*.

**Figure 4.**
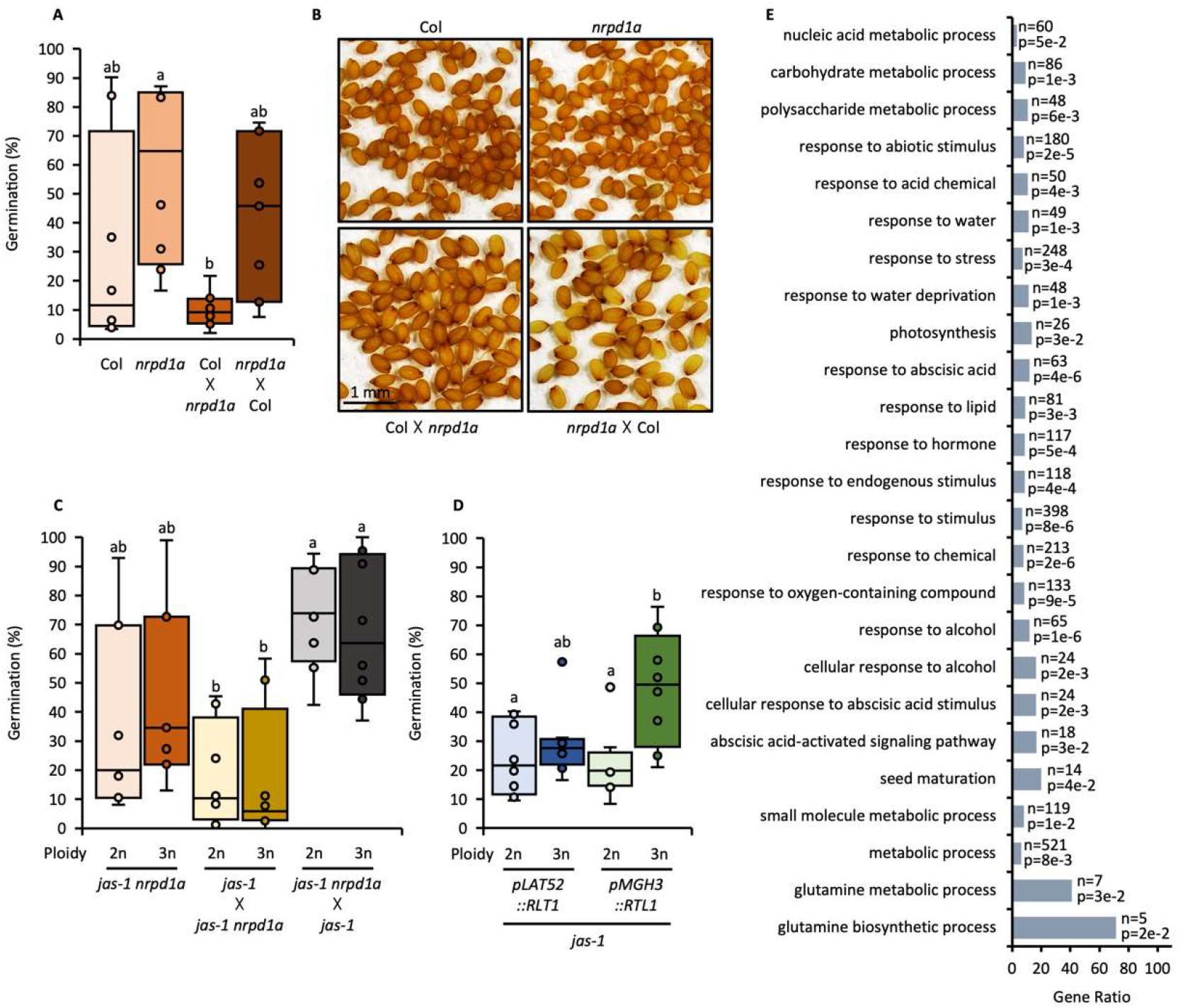
RNA PoIV-dependent processes control seed dormancy in polyploids with paternal genome excess. **A**. Germination of Col, *nrpd1a*, maternal Col-0 x *nrpd1a*, and *nrpd1a* x Col-0 seeds set at 15 °C and stratified for 3 days. Significant differences are shown using ANOVA with Tukey post hoc test (P < 0.01 n = 7-8). **B** Representative images of mature seeds set at 15 °C from Col, *nrpd1a* and reciprocal crosses. **C** Germination of *jas-1 nrpd1a* diploid (from diploid *jas-1 nrpd1a* plants), *jas-1 nrpd1a* triploid (from diploid *jas-1 nrpd1a* plants) seeds, and seeds from reciprocal crosses set at 15 °C and stratified for 3 days. Significant differences are shown using ANOVA with Tukey post hoc test (P < 0.01 n= 7-8). **D** Germination of *jas-1 pLAT52::RLT* diploid (from diploid *jas-1 pLAT52::RLT* plants), *jas-1 pLAT52::RLT* triploid (from diploid *jas-1 pLAT52::RLT* plants), *jas-1 pMGH3::RLT* diploid (from diploid *jas-1 pMGH3::RLT* plants), *jas-1 pMGH3::RLT* triploid (from diploid *jas-1 pMGH3::RLT* plants) seeds set at 15 °C and stratified for 3 days. Significant differences are shown using ANOVA with Tukey post hoc test (P < 0.05 n= 8). Boxplots indicate minimum and maximum values, as well as 25th, 50th and 75th percentiles. **E** GO-Term analysis of up-regulated genes in endosperm of nrpd1 x WT vs WT developing seeds (Satyaki and Gehring 2022), showing gene function category, p-values and number of genes in each category.

Paternal siRNAs affecting the triploid block originate in the vegetative cell (Pachamuthu et al., 2024). To further demonstrate the role of paternal siRNAs in dormancy reduction we compared jas-1 mutants crossed to expressing RNASE THREE-LIKE 1 (RTL1) in either the pollen vegetative or sperm cells (Figure 4D; Pachamuthu et al., 2024). RTL1 prevents siRNA production by cleaving doubled stranded RNA, (Shamandi et al., 2015). We found that *jas-1* diploid and triploid seeds showed identical dormancy levels when combined with the expression of RTL1 under the pLAST52 vegetative cell promoter. In contrast, dormancy differences remained when RTL1 was expressed under the sperm cell-specific MGH3 promoter (Figure 4D). This further confirmed that elimination of paternal siRNAs prevents paternal genome excess from blocking seed dormancy induction by the mother plant.

Maternal loss of *NRPD1* leads to mis-expression of large numbers of genes in the endosperm, including up-regulation of dormancy-imposing gene *DELAY OF GERMINATION 1* (Satyaki and Gehring, 2022). To understand dormancy-related gene expression in *nrpd1* x WT seeds we performed GO-term analysis of endosperm RNAseq data comparing *nrpd1* x WT endosperm to WT (Satyaki and Gehring 2022). This surprisingly showed that up-regulated genes included many involved in ABA-regulation of seed dormancy such as *AHG3, DOG1, DOGL4* and multiple ABA-responsive genes, suggesting that the ABA response of the endosperm is elevated compared to WT seeds (Figure 4E). This is inconsistent with the low dormancy phenotype, but consistent with our analysis of endosperm transcriptomes in seeds with paternal excess (Figure 3C). Interestingly, we also see increased expression of genes associated with inner integument functions such as cuticle synthesis and phenylpropanoid metabolism, a phenomenon was can also detect in seeds with paternal excess.

## Discussion

The activity of maternally and paternally derived genomes in the endosperm is often viewed through the prism of parental conflict, but most commonly over changes to resource acquisition caused by alternations in seed size (Scott et al., 1998). Here we show that the antagonistic activity of maternally and paternally derived PolIV-dependent RNAs during seed development is integral to conflicts in seed dormancy. Paternal genome dose lowers seed dormancy and some of our experiments also reveal a complementary antagonistic dormancy-promoting of the maternal genome. Our assays were not optimised to detected this latter effect but it is consistent with previous studies (Iwasaki et al., 2019; Sato et al., 2021; Chen et al., 2023) that reveal roles for the maternal epigenome in dormancy induction. These low dormancy states can be viewed as an attempt by the father to subvert maternal bet-hedging strategies which can lead to risky dormancy and dispersal strategies for individual progeny seeds within her seed lot (Penfield, 2017).

Studies comparing dormancy in naturally occurring diploids and polyploids of the same species in general show that polyploids are more likely to show higher levels of dormancy than their diploid relatives (Tyler et al., 1978; Stevens et al., 2020), although in these studies the effect of polyploidisation is confounded by genetic differences between varieties, which given the high selection pressure on seed dormancy traits (Huang et al., 2010) is likely to be substantial. Here comparing otherwise genetically near-identical polyploids with their diploid progenitors we show that polyploidisation can have a similar initial dormancy-reducing effect to increasing paternal genome dose alone (Figure 2). Thus in Arabidopsis the effect of the paternal genome is greater per unit dose than the maternal genome, which is interesting given that in the balanced state the endosperm has evolved to contain two copies of the maternal genome.

As shown by the phenotypes of *nrpd1* mutants, the effect of parental genome dose requires RNA PolIV-derived RNAs from both parents (Figure 4). Interestingly, paternal PolIV dose appears irrelevant in the absence of maternal PolIV-derived RNAs (Figure 4A) in a mirror image of the dominant role of the paternal genome in lowering seed dormancy (Figure 2. Thus the role of paternal PolIV-derived RNAs is to antagonise the dominant function of maternal RNAs in seeds. Interestingly, this role may not relate to endospermic 24-nt siRNA production and resulting RNA-directed DNA methylation because *nrpd1* mutants have normal levels of CHH DNA methylation in the endosperm (Cao et al., 2003; Iwasaki et al., 2019), despite a strong seed dormancy phenotype. However, these observations are consistent with observations in *Brassica rapa* that Pol IV loss in maternal sporophytic tissues results in high rates of post-germinative seed abortion (Grover et al., 2018). Interestingly, *B. rapa nrpd1* mutants also show seed coat colour alterations, a phenomenon we observe in Arabidopsis if only the mother is PolIV deffective (Figure 4B). Pollen vegetative cells predominantly accumulate so-called epigenetically-activated 21-22 nt siRNAs (easiRNAs) in a Pol IV-dependent manner, rather than 24 nt siRNAs that participate in RdDM (Martinez et al., 2018; Panda et al., 2020), but these have only limited effects on endospermic gene expression (Satyaki and Gehring, 2022), leaving their function in seed dormancy unclear. In contrast loss of maternal Pol IV-derived siRNAs leads to wide ranging effects on dormancy-associated gene expression with key transcripts relating to the induction of dormancy by ABA being up-regulated in maternal *nrpd1*/+ seeds (Satyaki and Gehring, 2022), despite the fact that these seeds have lower levels of dormancy (Figure 4A). So although parentally-derived siRNAs may play an important role in the control of progeny dormancy by parents, their mode and site of action remains unknown.

## Materials and Methods

### Plant materials and growth conditions

*Arabidopsis thaliana* mutants used in this study which have been previously described: *jas-1* (Erilova et al., 2009), *jas-3* (*jas*) (SAIL_813_H03; Erilova et al., 2009), *nrpd1a-3* (*nrpd1*) (SALK_128428; Herr et al., 2005), *jas nrpd1* (Pachamuthu et al., 2024), *jas-3 pLAT52::RLT1* (Pachamuthu et al., 2024), and *jas-3 pMGH3::RLT1* (Pachamuthu et al., 2024). Tetraploid Ler and *jas-1* were generated by colchicine treatment and stabilisation for XXXX generations. Tetraploid Col was ordered from Nottingham Arabidopsis Stock Centre (N69114). All mutants are in Col-0 background except *jas-1*, which is in the Landsberg (Ler) background, and *vel3-1* (SALK_052041; Chen et al., 2023) that was backcrossed 4 times into Ler (*vel3*^*bc*^). Arabidopsis plants were directly grown in long days (16h light and 8h darkness at 22°C) until bolting. For seed dormancy assays mother plants were moved to long days at 16°C for the duration of seed set to ensure strong dormancy for the experiments.

### Seed germination assays

For dormancy assays, freshly matured seeds were sowed onto 0.9% water agar plates and placed in long days (16h light and 8h darkness) at 22°C. Stratification was performed at 4°C in the dark for 3 days as indicated. For ABA sensitivity, seeds were sowed onto 0.9% water agar plates with or without exogenous ABA, stratified, and then placed in long days (16h light and 8h darkness) at 22°C. Seed batches from individual mother plants were used as biological replicates, at least 4 biological replicates of ∼50 seeds were used. Seed germination was scored by radicle emergence at 7 days after sowing or 4 days after stratification as indicated.

### ABA measurement

ABA were measured as previously described (Chen et al., 2023) using seed lots from individual mother plants as biological replicates.

### RNA-seq

RNA-seq was performed as described in Chen et al. (2023). In short, RNA was extracted from mature endosperm by first imbibing 0.05g of seeds for 1 hour. Following Iwasaki & Lopez-Molina (2021), endosperm-enriched tissues were then isolated using a glass slide to squeeze the seeds and centrifuged twice at maximum speed with 40% sucrose. After washing three times with water, the samples were stored at -80°C. RNA extraction followed the borate method (Penfield et al., 2005). RNA sequencing was performed by Novogene using a Hiseq 4000 system, generating 150 bp paired-end sequences with a minimum of 25 million reads per sample. Three biological replicates were prepared. Raw RNA-seq reads were trimmed using Cutadapt and mapped to the *Arabidopsis thaliana* TAIR10 genome with STAR. Read counts for each gene were obtained using featureCounts, and only sense reads were analyzed further. Differentially expressed genes (DEGs) were identified with edgeR, using FDR < 0.05 and a fold-change ≥ 2. For visualization, bigwig files were generated with deepTools and viewed in IGV.

